# Muscle Activation Profile During Perturbed Walking is Modulated According to Body State

**DOI:** 10.1101/2021.01.13.426393

**Authors:** Uri Rosenblum, Itshak Melzer, Gabi Zeilig, Meir Plotnik

## Abstract

During an unexpected loss of balance, avoiding a fall requires people to readjust their footing rapidly and effectively. We investigated the activation patterns of the ankle and knee muscles, and muscle fiber type recruitment resulting from unannounced, medio-lateral (i.e., right/left) horizontal-surface walking perturbations in twenty healthy adults (27.00±2.79 years, 10 females). Surface electromyography (sEMG) total spectral power for specific frequency bands (40-60Hz, 60-150Hz, 150-250Hz, 250-400Hz and 400-1000Hz), from tibialis anterior (TA) and vastus lateralis (VL) muscles were analyzed. Compared to non-perturbed walking, we found a significant increase in the total spectral power of lower-extremity muscles during the first 3 seconds after perturbation. When two feet were on the ground in time of perturbation we found a different muscle fiber type recruitment pattern between VL and TA muscles. This was not significant for perturbations implemented when one foot was on the ground. Our findings suggest that muscle operating frequency is modulated in real time to fit body state and somatosensory input in the context of functional goal requirements such as a rapid change of footing in response to unexpected loss of balance in single and double-support phases of gait.

**New & Noteworthy:** To study muscle spectral profiles in response to loss of balance, we investigated the dynamics of muscle spectral power changes, across different frequency bands after unannounced mechanical perturbations during walking. We showed increased activation of high-frequency motor units of the lower-limb muscles, subside 3 seconds after perturbation. Differences in power increase of specific frequency bands suggest that muscle activation is modulated in real time to fit body state in the context of functional goal requirements.

## Introduction

Stability while walking is maintained by utilizing two mechanisms: (1) *postural orientation*: active body alignment with respect to gravity, the supporting surface, and visual environment. It is achieved through constant reweighting of integrated somatosensory, visual and vestibular information by the central nervous system (CNS) (2) *postural equilibrium*: active control over the placement of the body center of mass (CoM) commensurate with its velocity (also known as the extrapolated CoM, XcoM) and within the boundaries of its base of support (BoS) provided by the feet (Hof, 2008a; Horak, 2006). The distance between the BoS and the XcoM is a measure of dynamic stability called margin of stability (MoS) (Hof et al., 2005). Thus, in response to external-or self-disturbances to stability, the CNS utilizes these mechanisms, by means of coordinating movement between arms, trunk and lower-limb muscles, to maintain equilibrium (Berger et al., 1984; Horak, 2006; Tang et al., 1998).

In laboratory settings, unexpected external perturbations are introduced in different phases of the gait cycle to provoke postural disturbances. To avoid falling, individuals perform rapid, reflex-like compensatory stepping responses to modulate their BoS (i.e., footing) by stepping in the direction of the CoM velocity (Bruijn and van Dieën, 2018; Hof et al., 2010). For example, when the CoM is displaced outward, usually an inward compensatory stepping response is performed to prevent fall (see Fig. S1 in *supplementary materials)*. Compensatory stepping responses to perturbations are accompanied by elevated sEMG activity of the distal (Mueller et al., 2018; Tang et al., 1998) and proximal (Tang et al., 1998) lower limb muscles (e.g., see Fig. 1 and video in *supplementary materials*) and suggested to be scaled in relation to the XcoM displacement magnitudes (Chvatal and Ting, 2013; Hof and Duysens, 2018). This compensatory stepping response was reported to be under spinal, supraspinal and cortical control (Bruijn and van Dieën, 2018; Jacobs and Horak, 2007). It is characterized by sequences of movement strategies (e.g., outward & inward stepping) and muscle activity can be measured using surface electromyography (sEMG) to describe muscle recruitment patterns and synergies (Berger et al., 1984; Chvatal and Ting, 2013; Hof et al., 2010; Hof and Duysens, 2018, 2013; Tang et al., 1998).

**Fig.1:**
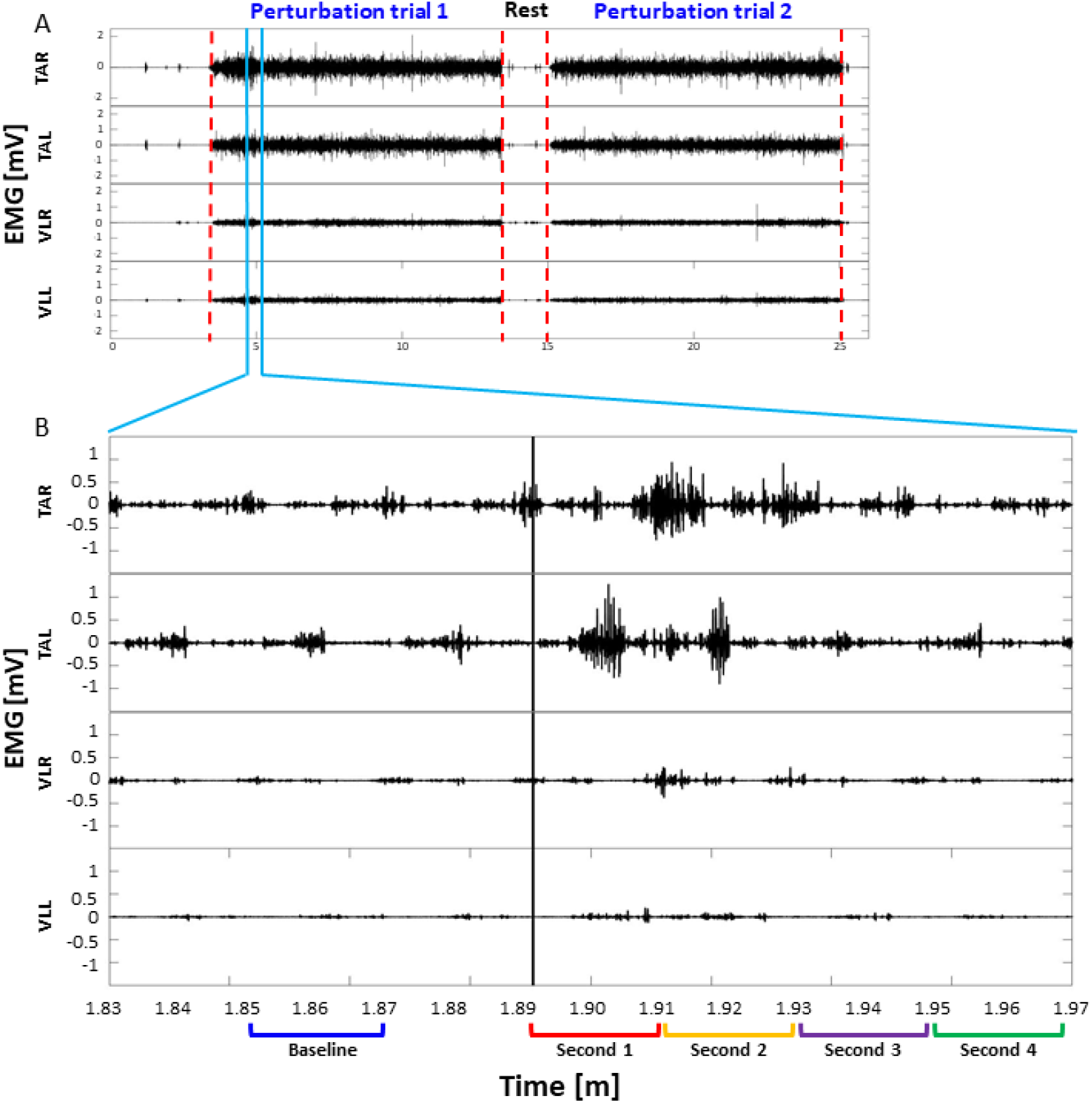
Filtered surface electromyography, sEMG. Data for 2 exemplary, consecutive trials (A) and for one specific perturbation in a window of 1 second bin, 2 seconds before (baseline) and after (second 1-4) the perturbation (B), of left and right tibialis anterior (TA) and vastus lateralis (VL). From the traces we can see that TA activity is greater than VL activity for the whole 2 trials and in the specific perturbation. Worth noting that the signals in panel A maintain relatively constant amplitude throughout the trials, indicating no effects of fatigue. Additionally, in panel B we can see the muscle response very evident in the first 2 seconds after perturbation onset (black vertical line) after which it gradually returns to pre-perturbation values.

The CNS has two general means to control the force exerted by skeletal muscles: (1) altering the number of active motor units, called recruitment; (2) modulate the rate of action-potential impulses (i.e., frequency) driving motor units, called rate coding. Motor units are recruited by size (i.e., *the size principle*) from those that produce the smallest forces (Type I) to those that exert the largest forces (Type II) (Fling et al., 2009; Henneman et al., 1965; Olson et al., 1968; Zajac and Faden, 1985). (Kupa et al., 1995; Wakeling et al., 2001). In humans, lower frequencies, i.e., 40-60Hz are attributed to the activity of small alpha-motoneurons (type I, slow-twitch muscle fibers), 60-120Hz are attributed to medium alpha-motoneurons (type IIa, fast-twitch muscle fibers), and 170-220 Hz, are attributed to the large alpha-motoneurons (type IIb, fast-twitch muscle fibers) (Wakeling et al., 2001). Frequencies of 400Hz and above are associated with the beginning of a ballistic or dynamic contractions and rapid force production (Christie and Kamen, 2006; Cutsem et al., 1998).

Recently, interest has grown in muscle activation frequency while walking (Chan et al., 2016; Hussain and Mamun, 2012; Roeder et al., 2020; Zhang et al., 2017). However, the dynamics of muscle spectral power density in response to unexpected balance loss during balance perturbations has not been investigated. Deeper understanding of muscle activation patterns in response to unexpected balance perturbations will provide insights to the mechanisms of balance recovery responses and could have implications for treatment of people with balance deficits such as the elderly or patients with neurodegenerative/neurological disorders.

We therefore sought to investigate the differences between responses to perturbations in single-support compared with double-support in terms of behavior (i.e., stepping and dynamic stability characteristics) and lower-limb muscle activation pattern, as measured by sEMG. Three hypotheses drive our work: (1) Total spectral power of lower limb muscles would increase due to unannounced perturbations while walking, then gradually decrease; (2) total spectral power of sEMG frequencies associated with type IIa and IIb muscle fibers will increase more rapidly than lower frequencies associated with Type I muscle fibers to provide rapid force exertion. We hypothesize this will be more prominent in single-support condition due to the greater demand from the stance limb during perturbation; (3) As TA muscles are strongly implicated in balance recovery while walking (Hof et al., 2010), and perform finer movements than VL (i.e., shifting the pressure under the foot to contain the body CoM), we hypothesized that the TA muscle will demonstrate greater change in higher frequency bands compared to VL muscle, similar to the results found by Seki and Narusawa (Seki and Narusawa, 1996) for hand muscles.

## Results

One hundred and sixty-five perturbations were analyzed (68 for single-support and 97 for double-support phases of gait cycle). All participants successfully recovered from the perturbations without falling. We did not see a habituation to the protocol as stepping times did not change significantly and muscle activation patterns remained consistent over the course of the experiment.

### Dynamic stability and stepping response to rightward surface translation, during left leading limb stance, while walking

Stepping response strategy was maintained over time across participants (see Fig. S2 in *supplementary materials*). All participants reacted with inward step to rightward surface translations occurring in left midstance or double-support with left leading limb, as predicted by the inverted pendulum model (Hof, 2008b; Hof et al., 2010, 2005). Behavior outcomes of stepping responses and dynamic stability in response to rightward surface translation, during left limb stance are summarized in Table 2 and characterized in detail in the Supplementary materials.

Fig. 2B-D summarizes the 1^st^ mixed-effect model (F_[11,966]_ = 19.22, p<0.001). The model showed a significant interaction effect of time by condition (F_[5, 966]_ = 4.84, p<0.001, Fig. 2B) as well as significant main effects for time segment (F_[5, 966]_ = 34.71, p<0.001, Fig. 2C) and condition (F_[1, 966]_ = 4.98, p=0.026, Fig. 2D). The post hoc analyses revealed significantly larger average MoS in the 1^st^ second after perturbation compared to all other strides (p<0.001), except for 2 seconds after perturbation onset (p=0.055), see Fig. 2C. The average MoS in the 1^st^ and 2^nd^ seconds after perturbation were significantly greater than MoS in baseline (p<0.001, Fig. 2C). In time of perturbation as well as in the 1^st^ and 3^rd^ second after perturbation onset, in single-support condition average MoS is significantly greater than double-support condition (p<0.05, Fig. 2B).

**Fig. 2:**
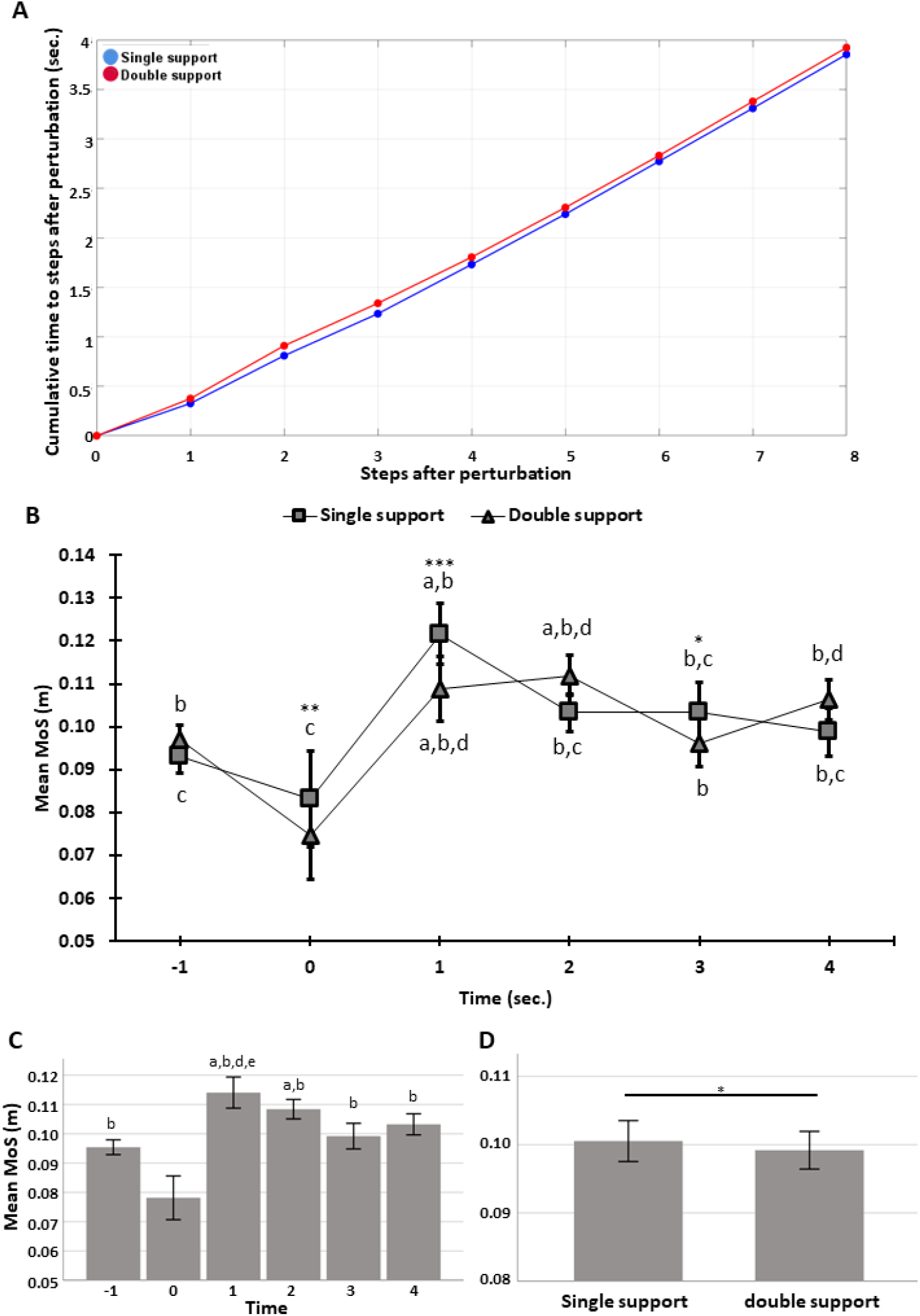
cumulative average stepping time after perturbation onset for all participants (n=20, A) and mixed-effect model for evaluating time and condition effect on dynamic stability using averaged margin of stability (MoS) before-at the instance- and after-the perturbation onset for all participants (n=20, B). In A and B blue traces represent single-support data and red traces are for double-support data. (A) X axis represents number of steps after perturbation onset. Y axis represents time in seconds. On average every two steps (i.e., a stride) are completed within a second; (B) X axis represents time in seconds, where 0 is the perturbation onset time, -1 is for 1 second (i.e., one stride) of baseline (before perturbation) and positive values are time after perturbation onset. Y axis represents average MoS in meters; (C) main effect of time in seconds, where 0 is the time of perturbation onset. (D) Main effect for condition (i.e., single-support vs. double support). Error bars represent 95%CI of the mean. a – significantly greater than baseline; b - significantly greater than Perturbation time; c – significantly less than MoS in stride 1; d – significantly greater than MoS in stride 3; e – significantly greater than MoS in stride 4; * - significant difference between single and double-support p<0.05. ** - significant difference between single and double-support p<0.01. *** - significant difference between single and double-support p<0.001.

### Spectral power distribution at baseline and four seconds after perturbation for all frequencies, muscles and conditions

Spectral power distribution for the left and right TA and VL before and after perturbations are presented in Fig. 3A-H. Since there are differences in the soft tissue thickness covering the TA and VL, total spectral power (i.e., area under curve of profiles in Fig.3A-H) was normalized to standing condition to allow for muscle activity comparison. The mixed-effect model (F_[39,1200]_ = 10.53, p<0.001) showed significant main effect for time segment (i.e., one second bin two seconds before the perturbation (i.e., baseline) to the 4^th^ second after perturbation; F_[4, 1200]_ = 22.28, p<0.001, Fig. 4A), and muscles (F_[3, 1200]_ = 99.96, p<0.001, Fig. 4B). More specifically, there was a significant increase in the total spectral power (S(f)) for all muscles combined in the 1^st^ three seconds after perturbation, as compared to baseline (p<0.02), such that the largest increase occurred during the first second after perturbation, followed by a gradual decrease until the fourth second. These results support our 1^st^ hypothesis that the total spectral power for all frequencies would increase due to unannounced support-surface perturbations, and then gradually decrease, regardless of the perturbation type. Muscle S(f) in second 4 was not significantly different than baseline (p=0.38). Muscles’ main effect showed significantly higher S(f) for Right-TA muscle compared to all other muscles (i.e., Left-TA, Left-VL and Right-VL), p<0.001. Left and right TA muscles increased their activity significantly more than Left and right VL, respectively, p<0.001. We did not find a main effect for condition (F_[1, 1200]_ = 0.11, p<0.74). We found a significant muscle by condition interaction effect (F_[3, 1200]_ = 4.12, p=0.006). For example, the muscles’ S(f) pattern was consistent for both conditions (i.e., single and double-support) with the main effect, displaying higher S(f) values for TA compared to VL for the left and right limbs (p≤0.01). Finally, we found significantly higher S(f) for Left-TA in single-support than double-support (p=0.034) and significantly higher S(f) for Left-VL in double-support compared to single-support (Fig. 4c; p=0.014).

**Fig. 3:**
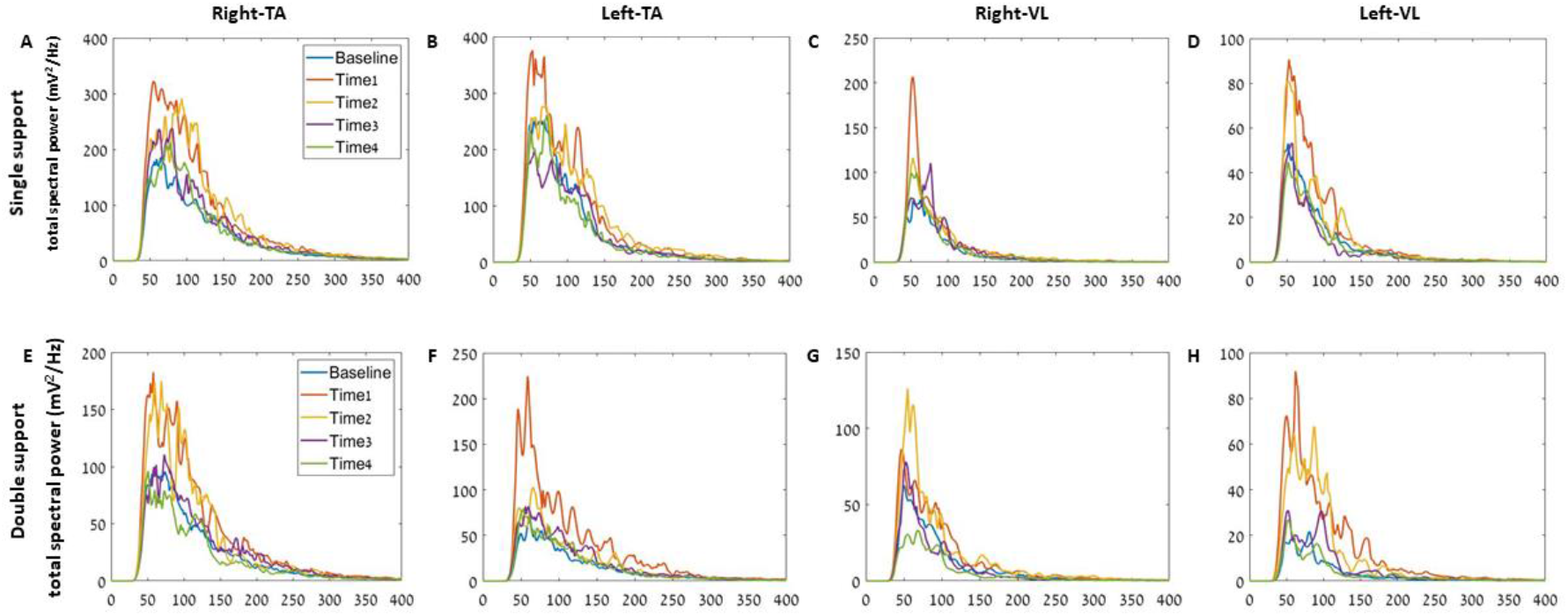
Progression of spectral power profiles over time in response to unexpected perturbations of rightward platform translation during single and double-support of the left leading limb. Ankle, right and left tibialis anterior (TA) muscle and knee, right and left vastus lateralis (VL) muscle (A, B, E, F and C, D, G, H, respectively) are depicted. Traces represent grand averages of spectral power density of raw surface electromyography (sEMG) data, across all participants (n=20). Please note that the second 3 and second 4 traces almost completely overlap with the baseline trace, indicating return to baseline.

**Fig. 4:**
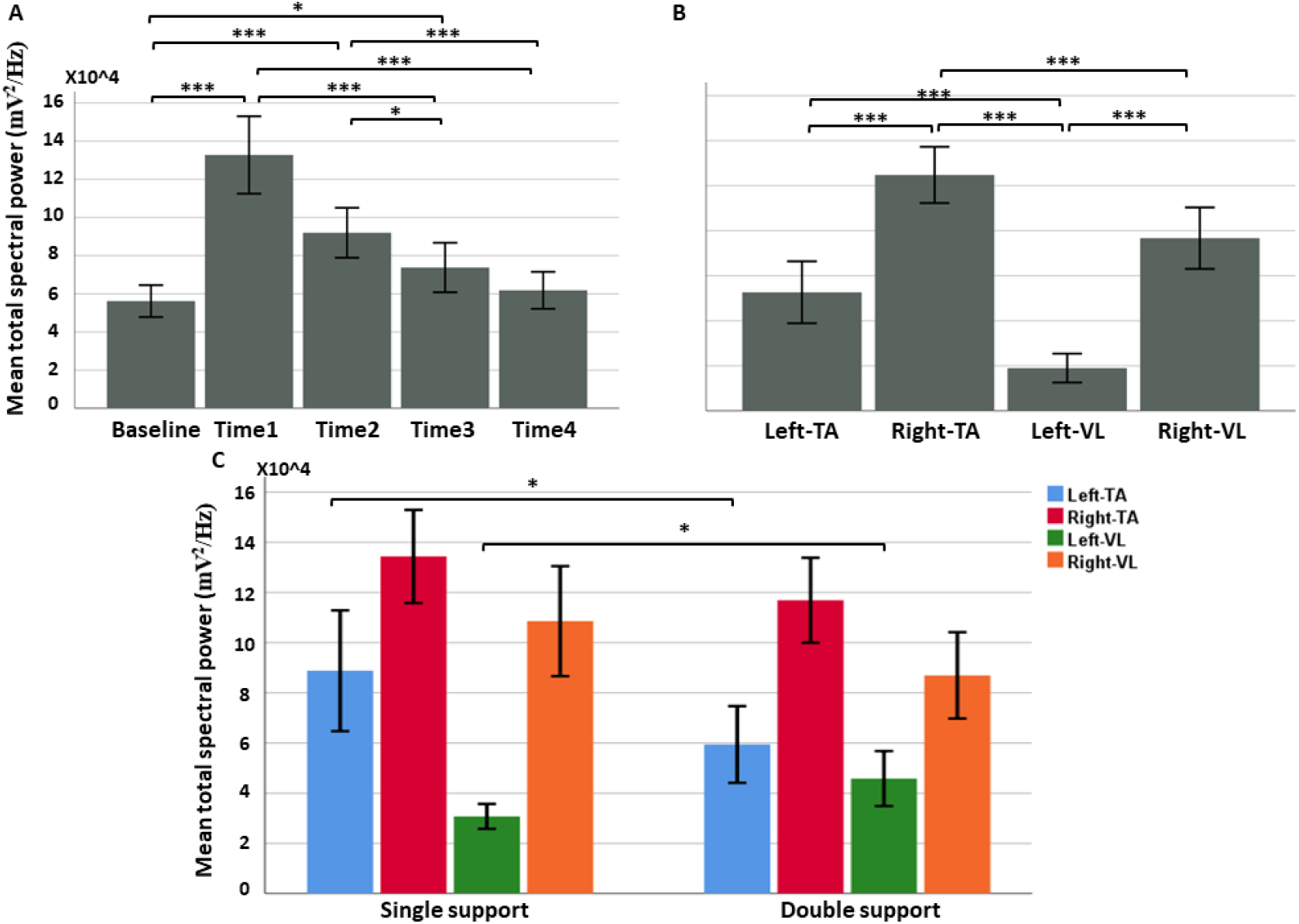
Summary of significant main effects and interaction of mixed-effects model to test our first hypothesis that total spectral power will increase after perturbation followed by gradual return to baseline values (n=20, df=1200). (A) time main effect, with post hoc analysis comparing mean total spectral power (S(f)) in baseline, represents 1 second (i.e., 1 strides) prior to perturbation onset, second 1-4 representing the 1^st^ to the 4^th^ second (i.e., 1^st^ stride until the 4^th^ stride) after perturbation; (B) muscle main effect, with post hoc analysis comparing S(f) for left and right tibialis anterior (TA) and vastus lateralis (VL); (C) interaction effect of condition by muscle showing higher Left-TA total spectral power in single-support compared to double-support and higher Left-VL values for double-support compared to single support. Values represent mean±95%CI. * P<0.05, ** P<0.01, *** p<0.001

### Temporal progression of leg muscle frequency band indicates muscle activation mechanisms are effected by body state

The mixed-effect model (F_[87, 4872]_ = 10.49, p<0.001) showed a significant main effects for time (F_[3, 4872]_ = 160.75, p<0.001). Significant change in total spectral power from baseline, from largest to smallest was second 1 > second 2 > second 3 > second 4, (p<0.001; see Fig. 5A), showing the result in the first model is robust. We also found a significant main effect for frequency band (F_[4, 4872]_= 2.63, p=0.033). 150-250 Hz exhibited significantly higher increase than 400-1000Hz (p=0.019). When we investigated the interaction of condition by frequency band (F_[4, 4872]_= 8.16, p<0.001) we found that in double-support perturbations there is a significant increase in all frequency bands compared with 400-1000Hz (p≤0.004, see Fig. 5C). No significant differences were found in single-support condition and double-support condition had greater increase of change from baseline for all frequency bands (p≤0.001). We also found that the Right-TA and Right-VL presented significantly higher increase in total spectral power change from baseline compared to Left-TA (significant main effect; F_[3, 4872]_= 5.75, p=0.001; p≤0.003). When investigating the interaction of muscle by condition (F_[3, 4872]_= 11.26, p<0.001) we found significantly higher increase in double-support condition compared to single-support condition for all muscles (p<0.001). In single-support condition we found the right (i.e., Right-TA/VL) increased their activity more than the left (i.e., Left-TA/VL) limb muscles, p≤0.001. In double-support condition we found the highest increase for the Left-VL compared to all other muscles (p≤0.018). Time by muscle by condition interaction (F_[18, 4872]_ = 1.67, p=0.037) showed the differences in muscle activation are attributed to second 2, where, in single-support Left-VL presented significantly lower increase than all other muscles (p≤0.023) and Right-TA did not present a significant increase. In double-support condition, Left-TA presented significantly greater increase than Left-VL and Right-VL (p≤0.035). In second 3, we only found significant differences in single-support where Left-TA had significantly lower increase than Right-TA and Right-VL (p≤0.045). No significant interaction effect was found for muscle by frequency band (F_[12, 4872]_ = 1.30, p=0.21). Finally, a main effect for perturbation condition (F_[1, 10,996]_= 220.40, p<0.001) showed a significantly greater increase for perturbations in double-support than in single-support (p<0.001). Every Main effect was tested for the other effects pooled together. A muscle by frequency band by condition interaction (F_[12, 4872]_ = 1.77, p=0.047, see Fig. 5E) reveled that in double-support condition the 40-60Hz band Left-VL exhibited greater elevation than left and right TA (p≤0.002) and Right-VL had also greater increase than Left-TA (p=0.002). In the 60-150Hz band Left-VL had significantly higher increase than all other muscles (p≤0.023). For Left-VL all frequency bands had greater increase than 400-1000Hz except for 250-400Hz band, p<0.001. 40-60Hz and 60-150Hz bands had greater increase than 250-400Hz. Right-VL exhibited significantly greater increase in 150-250Hz band than 400-1000Hz band (p=0.001). Note: Left-TA has a tendency for greater increase in 150-250Hz band than 40-60Hz band (p=0.086). No significant differences were found in single-support condition, however there was a trend towards higher increase in the 250-400Hz band in the right than the left limb muscles (p≤0.067). The interaction effect of time by frequency band by condition was not significant (F_[24, 4872]_ = 0.53, p=0.97).

**Fig. 5:**
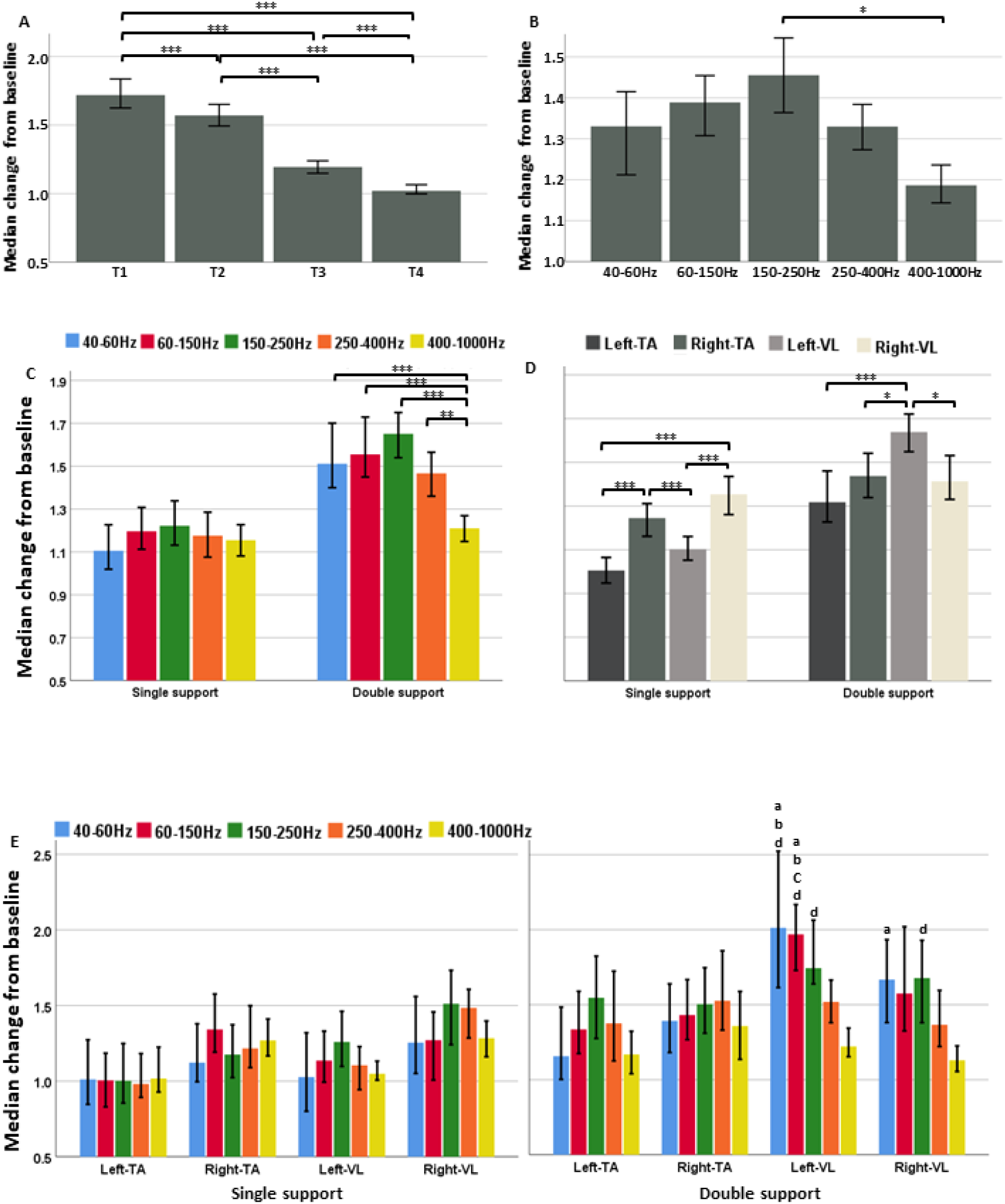
Summary of the 3^rd^ mixed-effect model of main effects and interactions relevant for our hypotheses testing that total spectral power, for all frequencies would increase due to unannounced support-surface perturbations, then gradually decrease, regardless of the perturbation type; total spectral power of frequencies associated with type IIa and IIb muscle fibers (i.e., 60-250 Hz frequency bands) will increase more rapidly than lower frequencies associated with Type I muscle fibers (i.e., 40-60 Hz frequency band); We further hypothesized this will be more prominent in single-support condition; TA muscle will demonstrate greater change in higher frequency bands compared to the VL muscle (n=4800); (A) main effect of time showing the dynamics in the change of frequency bands from baseline over time; (B) main effect of frequency band, showing highest increase for type II fibers related frequencies of 150-250 Hz followed by 60-150 Hz bands; (C) frequency band by condition interaction effect, showing the change in the different frequency bands is consistent between the conditions with different magnitude. Only changes in double-support condition are significant; (D) muscle by condition interaction effect, showing different muscle activation changes between double and single support; (E) muscle by frequency band by condition interaction effects. This effect showed a trend towards different muscle activation strategy between conditions. For all panels value of 1 represents no change from baseline, values less than 1 represent decrease relative to baseline and values greater than 1 represent increase activation relative to baseline. *p<0.05, **p<0.01, ***p<0.001, **a** = greater than Left-TA, **b** = greater than Right-TA, **c** = greater than Right-VL, **d** = greater than 400-1000Hz, **e** = greater than 250-400Hz.

## Discussion

As hypothesized, perturbation during single and double-support phases of gait caused a significant increase in the total spectral power in all frequency bands, followed by gradual decrease to baseline levels after ∼4 seconds (Figs. 4A and 5A). Our second hypothesis that S(f) of type II related muscles’ activation frequencies will increase more rapidly than type I related frequencies and that this will be more prominent in single support was not confirmed. We observed a significant difference in the increase in total spectral power in the 150-250Hz compared to 400-1000Hz frequency bands for the conditions combined. The interaction of frequency band by condition showed that the difference is driven by the double-support phase of gait cycle. When adding the muscle effect, we learn that specifically the left and right-VL muscles drive the difference above. This means that type IIb muscle fibers were more dominant in the response to perturbations in double-support phase than in single-support phase. Possibly, due to the smaller MoS in time of double-support perturbations (see Fig. 3B). Additionally, this is more pronounced in VL than TA muscles possibly due to the higher change in activation from baseline.

Our results further showed that, in the double-support phase, the recruitment of left-VL was greater compared to left-TA and right-TA for 40-60 Hz and 60-150Hz bands. Note, left-TA has a tendency for greater increase in 150-250Hz band than 40-60Hz band. These results refute our third hypothesis that the TA muscle will demonstrate greater change in high-frequency bands compared to the VL muscle. A possible explanation is that TA and VL differ in their architecture where as the tibialis anterior muscles contain about 70% slow type I fibers (Johnson et al., 1973), VL contain about 45-50% type I fibers (Johnson et al., 1973). Therefore, to activate type II muscle fibers to allow for rapid foot stepping we presume TA, a relatively small muscle compared to VL, is more code rating oriented while VL is more recruitment oriented. Furthermore, Previous work has shown that TA cross-sectional area of type I muscle fibers were smaller and type II muscle fibers larger for than for the VL, resulting in a larger type II/type I-muscle fiber cross-sectional area in the TA (Gregory et al., 2001).

### Total spectral power distribution and change over time for all frequencies combined

The observed increase in total spectral power for all frequencies in all muscles in the first second after unannounced perturbations (i.e., second 1) is in agreement with previous findings (Chvatal and Ting, 2013; Wakeling et al., 2001). This increase was followed by a gradual decrease of total spectral power to baseline levels, ending after 3 seconds (i.e., during second 3). The relatively prolonged elevated spectral power suggests the recovery process, which continues past the initial balance recovery response after perturbation (i.e., first 2-3 steps after perturbation (Hof et al., 2010; Lee et al., 2017; Zadravec et al., 2017)), is accompanied by increased muscle activation. The extended recovery process, beyond the initial response, is corroborated by kinematic outcomes of step length and width, as was described previously by us in details (Rosenblum et al., 2020), CoM displacement as was described by Ravi et al. (2021) and dynamic stability measures (i.e., MoS, see Fig. 3B-C).

### The role of muscle fibers sub types in responses to perturbations

Muscle activation spectral power signatures provide information on the muscles’ functional roles in locomotion (i.e., baseline walking in our study) and during recovery processes. Our results confirm that there is a modulation of fiber type activation of muscles in response to unannounced walking perturbations. Type II muscle fibers of the left and right TA and VL muscles were engaged as part of the rapid balance recovery response. This is expressed by the elevation in relevant frequencies (i.e., 40-60Hz and 60-150Hz band) in response to the perturbations, particularly in left-VL. Interestingly, our results suggest mirrored muscle recruitment strategy for balance response to single- or double-support perturbations, yet these effects are not significant (see Fig.5E). This could be explained by the behavior where in single-support condition there is a significantly larger MoS in second 1 compared to the double-support phase while in second 2 we see a significant decrease in MoS in single-support condition while in double-support we see a non-significant increase in MoS (see Fig.3B). This difference suggests the mechanism for muscle fibers’ recruitment pattern is not function specific (i.e., the same change in MoS from the time of perturbation to the stride after perturbation is achieved in the two conditions) but is influenced by body state estimation (i.e., lower MoS in time of perturbation in double-support).

### Differential effect between muscles

At the physiological level, the engagement of muscle activation for balance recovery was relatively greater for TA than VL in the same limb (see Fig 4B). We postulate that the muscles’ size difference along with their required function while walking and in response to perturbation explains this difference. The TA muscle fiber size is much smaller than that of VL. As a result, during dynamic movement such as walking and balance reactions to perturbations the extra load on the joints drives it to exert higher total spectral power. In response to unannounced perturbations, VL must maintain postural orientation, preventing the knee joint from collapsing under the body weight. The ankle muscles are mainly utilized for postural equilibrium, controlling ankle joint movements to redistribute ground pressure under the sole of the foot, effectively shifting the BoS and reducing the magnitude of a balance recovery step (Hof et al., 2010).

### Leg muscles fiber type recruitment relative to baseline in response to unexpected perturbations and progression over time

Our data indicate that type IIa muscle fibers contribute more to the increase in total spectral power than other muscle fiber types. Furthermore, our data suggests that the different muscles exhibit different muscle fiber type recruitment patterns in response to the perturbations in different conditions (see Fig.5E). This result is not in agreement with previous literature describing the size principle (Fling et al., 2009; Papagiannis et al., 2019). Other studies however, in animals and humans show that slow and fast muscle fiber recruitment pattern is essential for achieving optimal muscle performance to support effective whole limb movement (Holt et al., 2014; Wakeling et al., 2011). In our study, the participants are required to respond with accurate timing and precision to regain their balance. The trend towards different activation patterns of TA and VL muscles in the different conditions suggests different demands from the musculoskeletal system as dependent on the body state (i.e., single or double support) and not the functional demands of the task (i.e., taking a recovery step).

Findings described herein should be considered in the context of several limitations. First, our sample of 20 participants is rather small. A larger sample might have provided greater statistical power to show more robust differences in the muscle activation patterns. Second, it is worth mentioning that perturbations were relatively weak and slow compared, for example, to (Handelzalts et al., 2019; Nachmani et al., 2020). This might have prolonged the recovery process. Third, since only young adults were examined, the results may not be generalizable to middle-aged or older populations. Thus, future studis should be conducted to characterize the time-frequency domain of sEMG signals as well as the contribution of muscle fiber composition in relation to responding to unexpected walking perturbations in other populations (e.g., older adults, CVA, Parkinson’s disease). Lastly, this study only examined anterior lower limb muscles. Future studies including additional muscles (e.g., glutei) would provide a more complete picture of muscle activation patterns in response to unannounced perturbations during walking.

In conclusion, we were able to describe the temporal dynamics of muscle activation patterns in lower limb muscles. Our results revealed differential muscle activation patterns in response to different types of unexpected walking perturbations. Based on muscle activation patterns we show for the first time that the recovery process does not end after 2-3 steps, as was previously assumed. Our sEMG measurements demonstrate a residual balance recovery effect up to three seconds after perturbation. The pronounced change in Type II muscle fibers activation in the strides after perturbation and the trend to differences in muscle fibers’ recruitment between TA and VL muscles of the stance limb (i.e., left limb in our study) suggests different mechanisms of muscle activation in response to walking perturbations in different conditions (see Fig. 5E), in accord with the body state. In addition, our results support the notion that type IIa, and to some extent type IIb muscle fibers, are recruited with type I muscle fibers for rapid and effective balance recovery.

## Methods

### Participants

A convenient sample of 20 young adults, age 27±2.8, 10 females, participated in this study (Table 1). Since there are no sEMG frequency analysis studies in walking with perturbations, the sample size was chosen to be comparable with sEMG frequency analysis studies during walking with no perturbations using similar outcomes (Nüesch et al., 2012; Roeder et al., 2020; Zhang et al., 2017). Exclusion criteria included conditions effecting balance control such as: age over 65 (L. Sturnieks et al., 2008); obesity (i.e., body mass index [kg/m^2^] > 30) (Kopelman, 2000); orthopedic conditions affecting gait and balance (e.g., total knee replacement, total hip replacement, ankle sprain, limb fracture in the period of one year prior to participation (Jansen et al., 2013; Kubota et al., 2013, 2012; van Hoeve et al., 2019)); neurological diseases associated with loss of balance (e.g., multiple sclerosis, myelopathy); and conditions that could affect adherence and participation such as: cognitive loss or psychiatric conditions; untreated cardiac conditions (e.g., non-stable ischemic heart disease, congestive heart failure); and chronic obstructive pulmonary disease. The study protocol was approved by the Institutional Review Board of Sheba Medical Center). All participants provided written informed consent prior to entering the study.

**Table 1:**
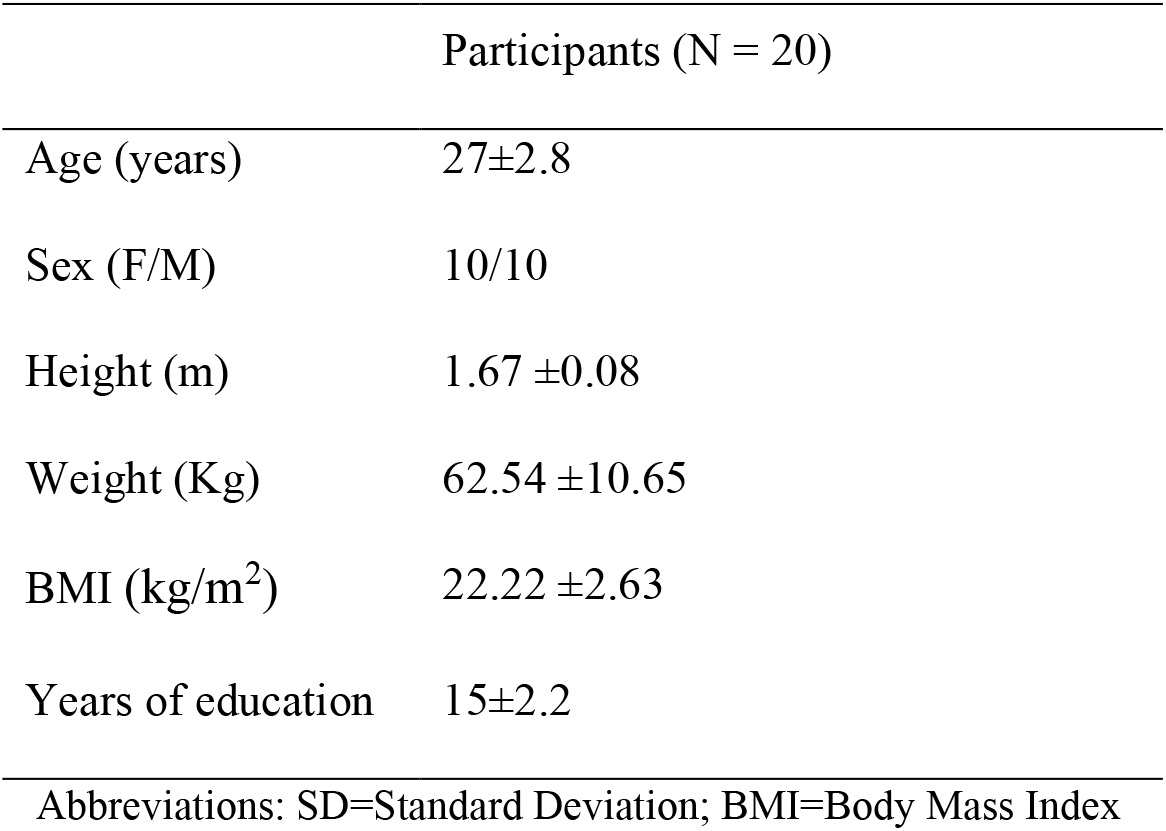
Demographic and physical characteristics (Mean ±SD)

**Table 2:**
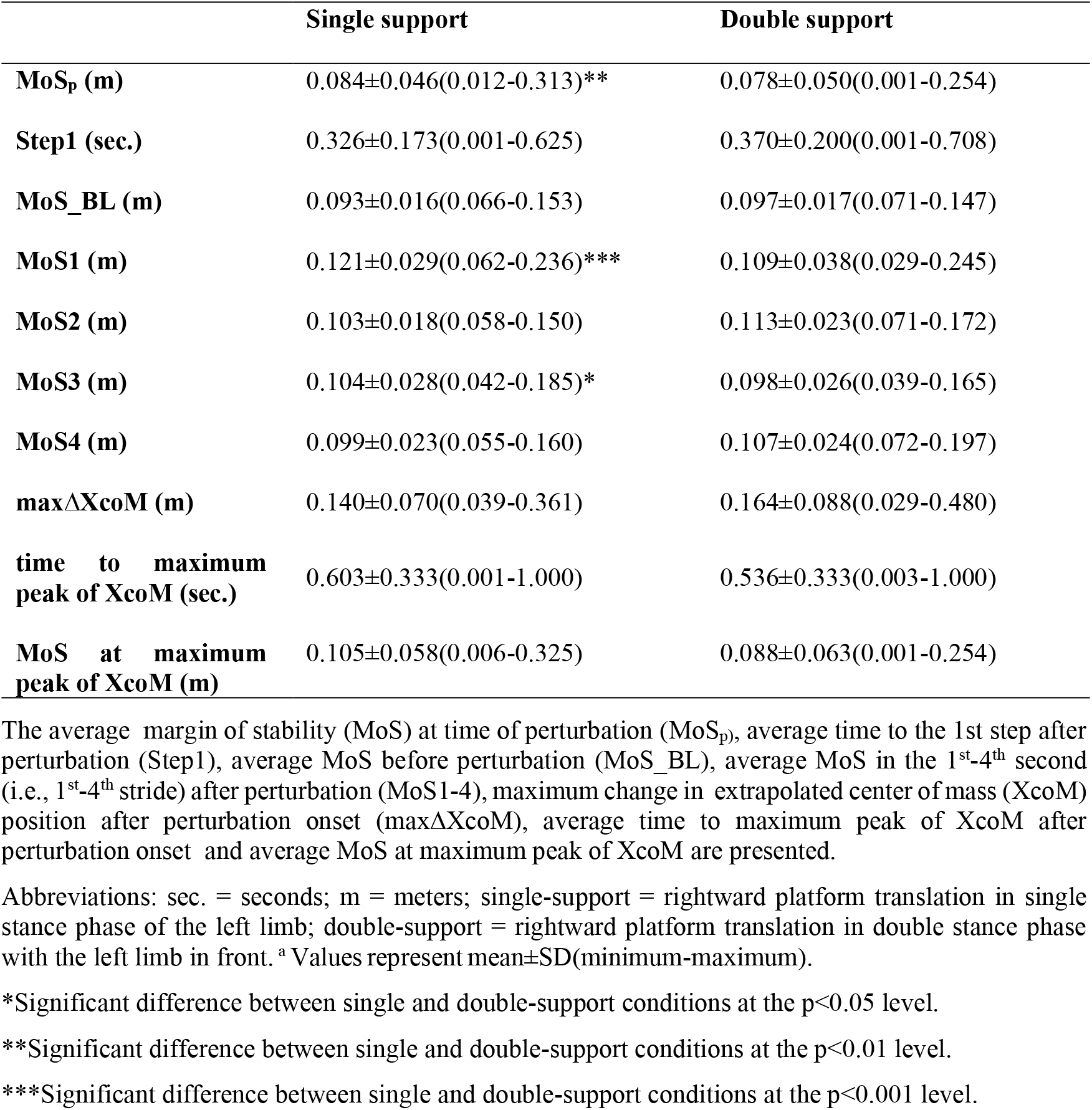
MoS and XcoM^a^ values in response to perturbations in single or double-support of the left leading limb.

### Apparatus and settings

Set-up for the experiment was described elsewhere in detail (Rosenblum et al., 2020). Briefly, participants walked in the CAREN High-End (Motek Medical B.V., Amsterdam, Netherlands) virtual reality system, equipped with a split-belt treadmill mounted on a platform that can be moved in six degrees of freedom. The platform is positioned in the center of a 360° dome-shaped screen. Unannounced support-surface perturbations along the anterior-posterior plane occurred by reducing the speed of one belt (split-belt treadmill) by 1.2 m/s with a deceleration of 5 m/s^2^. Right-left unannounced support-surface perturbations occurred by shifting the platform 15cm for 0.92 seconds, either left or right. A magnitude which induces changes in the gait pattern was chosen and was used for all participants for the sake of uniformity. Eighteen Vicon (Vicon Motion Systems, Oxford, UK) motion capture cameras placed around the dome’s circumference along with force plates under the treadmill’s belts, allowed for continuous recording of kinematic (motion) and kinetic (ground reaction forces) data, for real-time/post-hoc gait phase detection. Perturbation control and data recording were done by two computers that integrated kinetic, kinematic and perturbation data (see video in *supplementary materials*).

### sEMG recording

Raw sEMG data were recorded on a tablet computer (Lenovo ThinkPad) at 2048 Hz using an eego referential amplifier (eemagine Medical Imaging Solutions GmbH, Berlin, Germany), with an input impedance (referential) >1GΩ, and weight < 500g. Data were collected from left and right vastus lateralis (VL) and tibialis anterior (TA) muscles, known to take part in postural orientation (VL) and equilibrium (TA) (Chan et al., 2016; Hof and Duysens, 2018; Mueller et al., 2018; Papagiannis et al., 2019; Tang et al., 1998), as well as having a significant role in walking balance performance (Chvatal and Ting, 2013; Hof and Duysens, 2018; Nazifi et al., 2017). Additionally, there is reach literature on fiber type distribution for these muscles (Lexell, 1995; Staron et al., 2000, 1999; Zierath and Hawley, 2004). sEMG signals were recorded using bipolar hydrogel surface electrodes, 10mm in diameter, 24mm apart, placed parallel to muscle fibers, consistent with SENIAM guidelines (Hermens et al., 2000). The ground electrode was placed on spinous process of C7. Before electrode application, participants’ skin was shaved and cleaned with an abrasive pad and alcohol to reduce skin impedance. Finally, electrodes and cables were secured to the skin with tape to reduce moving artifacts. sEMG signals were tested for specific movements to control for cross-talk effects. Ankle dorsi/plantar flexion to assess TA activation and knee flex/extension to assess VL activation. sEMG data and perturbation events, kinetic and kinematic data were synchronized by the main, CAREN High-End, computer that started the Vicon system and provided a trigger to the eego amplifier once the virtual reality scene was activated.

### Protocol

The protocol consisted of three parts: acclimation to self-paced walking trial, static standing for 1 minute to serve as baseline for the sEMG activity and 6 walking with perturbation trials (48 perturbations for each participant). Between trials participants rested 2 minutes to avoid fatigue (see Fig. 1 for illustration; Note: no signs of fatigue were present in any of the participants’ EMG recordings, data not showed for the whole group). Participants first acclimated to walking in virtual reality environment (see (Rosenblum et al., 2020) for more details). During the acclimation phase they were instructed verbally by a physical therapist (UR) on how to operate the self-paced walking mode followed by a practice. In the practice they were asked to start walking slowly, rich their preferred walking speed and maintain it. The practice consisted of five consecutive walking trials with no perturbations, 30-45 seconds in each trial. In the second phase they were asked to stand with their hands at their sides, look straight forward at a plus sign that was projected on the screen in front of them and sway as little as possible for the duration of one minute. In the third phase the participants walked at a comfortable, self-selected speed (using the system’s self-paced mode (Plotnik et al., 2015)) while being subjected to random, unannounced perturbations at single or double-support phases of gait cycle and in different directions (anterior-posterior/medio-lateral), to minimize anticipatory responses and to provide a more ecologically valid environment. Note: for this paper only rightward, i.e., inward, platform translation perturbations in left single-support or double-support with the left leading limb were considered due to their relevance to falls and injury mechanisms in older adults’ population (Maki and McIlroy, 1997; McIlroy and Maki, 1996; Parkkari et al., 1999; Singer et al., 2016). Inward mediolateral surface translations elicit an inward stepping response, (see Fig S1 in *supplementary materials*) that is challenging for older adults’ population that show greater instability in midio-lateral direction compared to anterio-posterior direction (Maki and McIlroy, 1997; McIlroy and Maki, 1996; Singer et al., 2016). Furthermore, most hip fractures are caused by falling sideways (Parkkari et al., 1999). On the upside, these perturbations were shown to improve walking balance performance in older age as well as orthopedic and neurological conditions (Fitzgerald et al., 2000; Handelzalts et al., 2019; Lurie et al., 2020, 2013). Time intervals between consecutive perturbations was randomly varied (i.e., 25-35 seconds) to further minimize anticipation effect.

### sEMG data handling, processing, and analyses

sEMG data were preprocessed and analyzed across frequency domains using customized scripts written in MATLAB version R2019b (Mathworks, Nathick USA). Raw sEMG signals were first band-pass filtered (40–1000 Hz) (Papagiannis et al., 2019) with a 500^th^ order Butterworth filter to avoid movement artifacts found mainly in the lower frequency domain < 20 Hz (de Luca et al., 2010). Filtered sEMG signals were sliced in 1-minute windows around perturbation events (i.e., 30 sec. prior to 30 sec. after perturbation onset). sEMG data was considered for 1 second bin, 2 seconds prior to perturbation to serve as baseline, undisturbed walking data (see Fig.1B). One second bins, from 1-4 seconds after perturbation (second 1-4, respectively) served as balance and walking recovery sEMG data after perturbation. These segments were selected consistent with existing research from our group (Rosenblum et al., 2020) and Ravi et al. (2021) showing that most people recover gait within 4-6 seconds after perturbation onset during walking. We used one second bins, consistent with one stride (see Fig. 4A), therefore the terms stride and Time will be interchangeable. Perturbations were classified according to the leading limb and the platform translation direction (i.e., inward or outward perturbation). Since mediolateral perturbations are more challenging for the balance system (Osoba et al., 2019), only mediolateral perturbations were analyzed.

sEMG data processing consisted of two main steps following the procedures described by Garcia-Retortillo et al. (Garcia-Retortillo et al., 2020). In the first step, we quantified total spectral power, S(f), for averaged activity at baseline and second 1-4, for each muscle (i.e., right and left TA, and VL), by calculating the area under curves in Fig.3A-H. We used the ‘pwelch’ built-in MATLAB function to extract spectral power for each muscle and for every walking segment (i.e., baseline & second 1-4), using a time window of 100ms, with an overlap of 50ms. For each time window, we calculated total spectral power (S(f)) across all frequencies (Garcia-Retortillo et al., 2020):

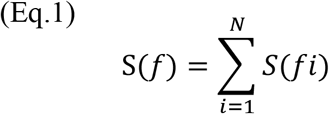

Where *fi* are all frequencies in the analysis, and N is the number of discrete frequency values for each segment.

To quantify differences in spectral power between segments, we calculated median total spectral power for each muscle, by calculating the area under the curves of Fig. 3A-H. Then the total spectral power of all time points was normalized to the total spectral power of the standing baseline trial to account for different morphological characteristics of the body around the muscles being measured (e.g., differences in body composition). In the second step, we measured the sequence in total spectral power over time from baseline to second 4 for each segment (see Fig. 3). We then examined the frequency spectrum of each segment in the following bands: 40-60Hz, 60-150Hz, 150-250Hz, 250-400Hz, and 400-1000Hz, based on (Garcia-Retortillo et al., 2020) and our data, presented in Fig. 3 and the progression of each frequency band in time (from baseline to second 4). Average spectral power (due to the different frequency band width) for every frequency band, in every segment was calculated using the following (Garcia-Retortillo et al., 2020):

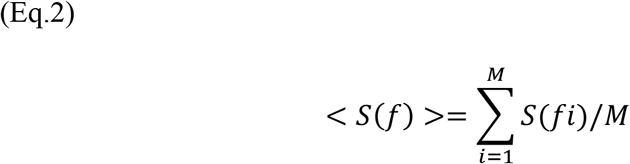

Where *f*_*i*_ are all frequencies in every frequency band, and M is the number of spectral power values in that frequency band. To specifically measure rate and magnitude of change across different frequency bands over time, we calculated change from baseline, for every frequency band, in every segment where, *fb* is the frequency band in every segment *x* (second 1-4) and *bl* is for the baseline segment:

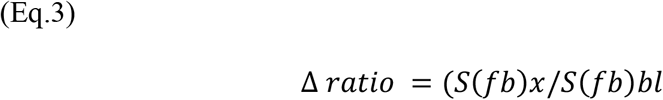

### Quantification of extrapolated center of mass and dynamic stability control

The margin of stability as a criterion for the state of dynamic stability during walking was calculated according to Hof et al. (Hof et al., 2005):

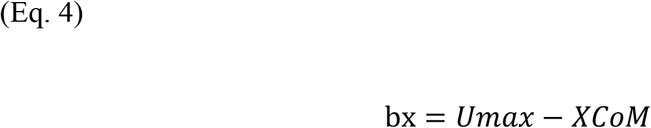

where b_x_ indicates the margin of stability in the medio-lateral direction, Umax is the lateral boundary of the base of support and XcoM is the position of the extrapolated center of mass in the medio-lateral direction (XcoM = P_XcoM_ + (V_CoM_/√(g/l))). P_XcoM_ is the horizontal (mediolateral) component of the projection of the center of mass to the ground, V_CoM_ is the horizontal CoM-velocity and the term √(g/l) presents the Eigen frequency of an inverted pendulum system with a leg of length l. Margin of stability was determined in the mediolateral direction since the extrapolated center of mass mainly shifts in mediolateral direction in response to mediolateral perturbations. the model is used to quantify dynamic stability during walking and can predict stepping reactions in response to shifts in XcoM due to perturbations. For each trial we identified: (a) initial contact of the disturbed leg (before experiencing the perturbation), (b) initial contact time of the recovery leg as well as the time to the next 7 steps after the perturbation, (c) the first local maximum of the mediolateral XcoM after the perturbation, (d) the MoS in mediolateral direction at time of perturbation, (e) MoS in mediolateral direction in time of the first local maximum of the mediolateral XcoM after the perturbation. (f) the average XcoM in every second after the perturbation, and (g) the average MoS in mediolateral direction in every second after the perturbation.

### Statistical analyses

To quantify and assess differences in the effect of condition (i.e., perturbations in single vs. double support) on dynamic stability (i.e., MoS) through the recovery process from unannounced perturbation while walking we computed mixed-effect models (1^st^ mixed-effect model) to account for dependence of observations within participants. MoS was found to have a normal distribution curve and was used as independent variable, while time segments (i.e., baseline through second 4) and condition along with their interaction (time segment by condition) were used as within-subject variables. To assess the difference in the change of MoS from time of perturbation to second 1 between single and double-support conditions, a paired T-Test was used. To test our first hypothesis (between-segment spectral power comparisons, across frequencies), we computed a 2^nd^ mixed-effect models with repeated measures to account for dependence of observations within participants. Since data did not distribute normally values were Ln transformed to fit a normal distribution. The transformed change was used as independent variables while time segments, muscles and condition (i.e., single/double support) along with the interactions of muscles by time segment, time segments by condition, muscles by condition and time segment by muscles by condition functioned as within-subject variables. To test our 2^nd^ and 3^rd^ hypotheses (between-segment spectral power comparisons, across frequencies, condition effect on total spectral power, differences in change in frequency bands across segments over time, TA muscle will demonstrate greater change in high frequency bands compared to the VL muscle) and to analyze total spectral power progression across frequency bands, we computed a 3^rd^ mixed-effect models with repeated measures to account for dependence of observations within participants. In each frequency band, we calculated change in total spectral power over time. Since data did not distribute normally, they were Ln transformed to fit a normal distribution curve. The transformed change was used as independent variable while time segments, frequency bands and muscles along with the interactions of frequency band by condition, muscles by condition, muscle by frequency band, muscle by frequency band by condition, muscle by time segment by condition, and frequency band by time segment by condition functioned as within-subject variables in mixed-effect models. Statistical significance was set *a priori* at p<0.05. Bonferroni correction was applied to account for multiple post-hoc comparisons. Data were analyzed using MATLAB version R2019b and SPSS version 23 (IBM Inc., Chicago, IL).

## Acknowledgements

The authors would like to thank Ofer Yizhar for the review of the manuscript. This study was supported in part by funding from the Israeli Ministry of Science and Technology, grant #3-12072, and from the Israel Science Fund, grant #3-14527. The research is part of one contributor’s (UR) work towards a doctoral degree from Ben-Gurion University of the Negev and was partially supported by a stipend.

## Supplementary materials

### Stepping behavior in response to walking perturbations

participants reacted with an inward 1^st^ step of the right leg in all (100%) instances, regardless of being in single or double support, as expected (see Fig. S1 in the *Supplementary martials*). Of the 2^nd^ steps, 37.32% and 76.04% were inward for single and double support, respectively, depending on the spatial constraints but not the step before. The 3^rd^ step was outward 100% of the times regardless of the 2^nd^ step direction and perturbation type (see Fig. S2 in *supplementary materials*). In other words, the participants’ CoM was accelerated laterally because of the medially moving platform, effectively disturbing XcoM trajectory (see Table 2 in main text for maximum change in XcoM and time to maximum peak of XcoM values). To avoid falling they must adjust their MoS, by performing an inward step (see Fig. S1 in *supplementary materials*), to insure the XcoM remains within the boundaries of the base of support (see the difference between MoS at time of perturbation and MoS at maximum peak of XcoM in Table 2). In the 2^nd^ step there are further corrections, inward or outward as needed, taking the space constraints of the treadmill borders into consideration. The 3^rd^ step is used to return to the center of the treadmill but still part of the recovery process as can be seen from the average MoS in the 2^nd^ stride (i.e., second 2, see Fig. 2B-C in main text) after perturbation (the average is still higher than the MoS before the perturbation, baseline values are attained in the 4^th^ stride, i.e., second 4, after perturbation, see Table 2 and Fig. 2B-C in main text).

**Fig S1:**
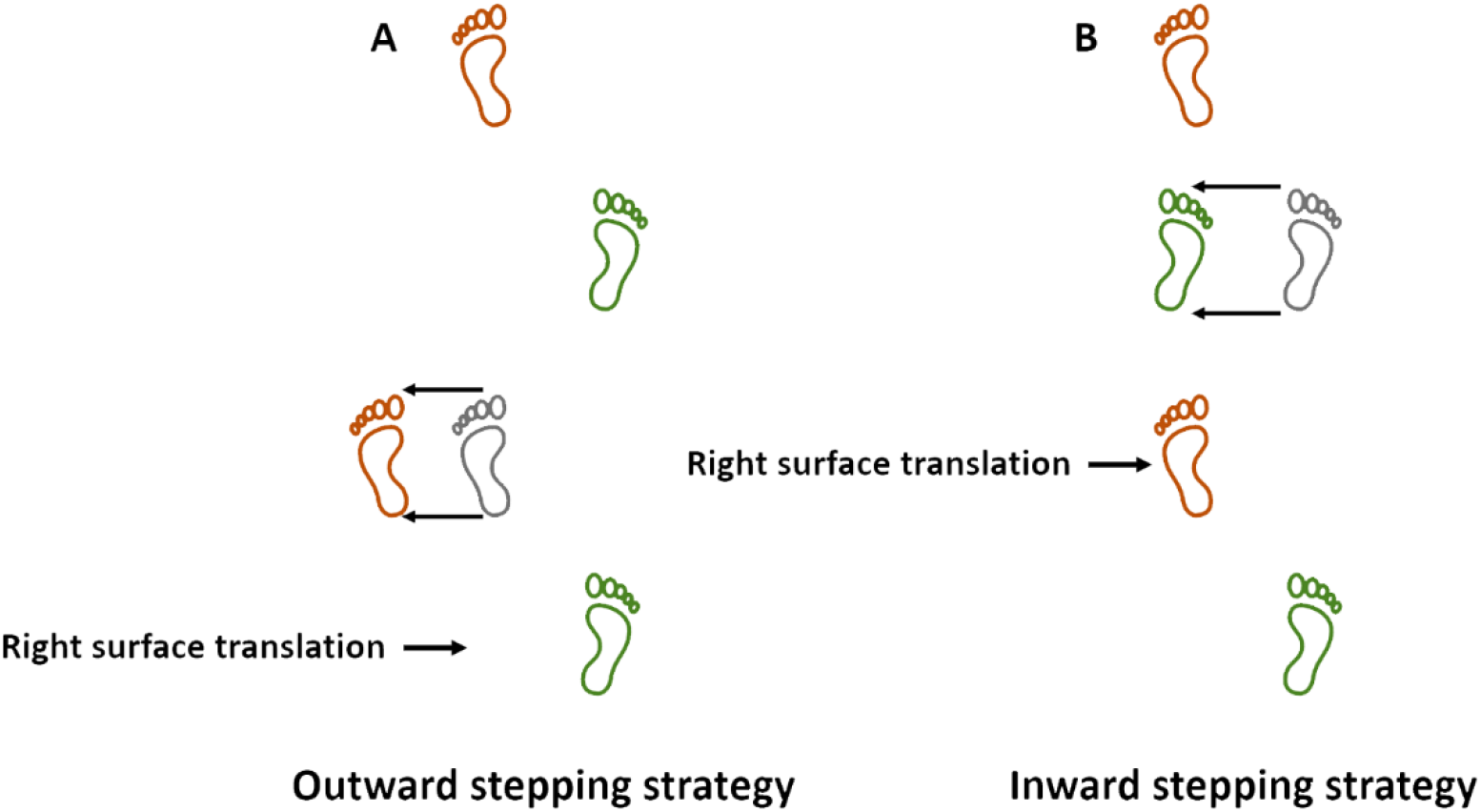
Stepping strategies in response to unexpected perturbations during walking. When exposed to rightward surface translation in right stance (A, green bottom foot) an outward stepping strategy with the left foot (orange) is required to maintain the center of mass in the boundaries of the base of support. Likewise, when exposed to rightward surface translation in left stance (B, orange bottom foot) an inward stepping strategy with the right foot (green) is required to maintain the center of mass in the boundaries of the base of support. If the right limb crosses the line of progression of the left limb, we call that a cross-over stepping strategy which is a specific case of the inward strategy.

**Fig.S2:**
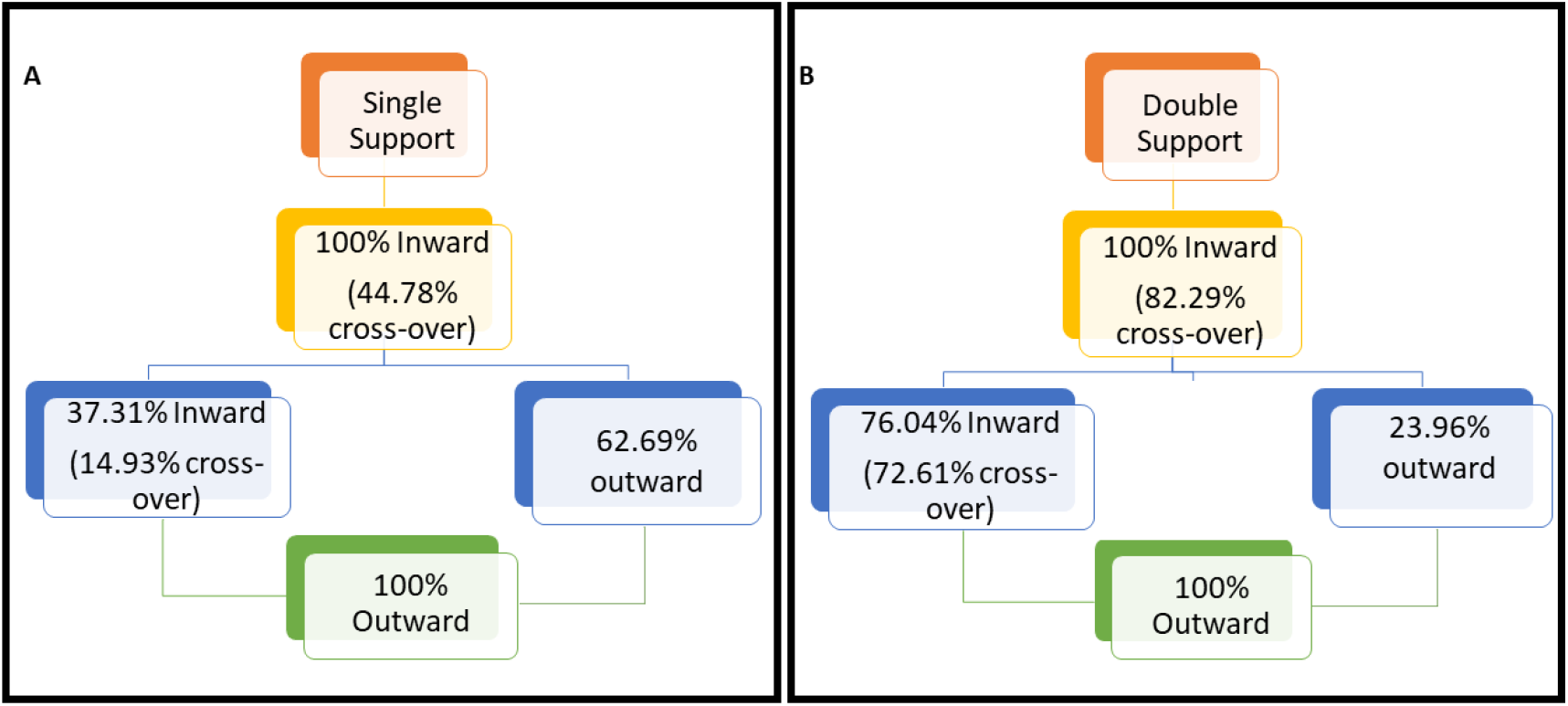
stepping reaction distribution in response to perturbations in single (A) and double (B) support conditions.

